# DEPDC1/ EEF1A1 complex promotes the progression of human osteosarcoma via downregulation of FOXO3a

**DOI:** 10.1101/2021.04.14.439766

**Authors:** Lin Shen, Han Li, Aijun Zhang, Ronghan Liu, Chendan Zhou, Ying Zhang, Kai Zhao, Morgan Bretches, Laitong Lu, Shang-You Yang, Bin Ning

**Affiliations:** Department of Orthopaedics, Jinan Central Hospital, Shandong First Medical University & Shandong Academy of Medical Sciences, Jinan, Shandong, 250013, China; Department of Endocrinology, The First Affiliated Hospital of Xiamen University, Xiamen University, Xiamen, Fujian, 350203, China; Department of Pediatrics, Qilu Hospital, Shandong University, Jinan, China; Department of Orthopaedic Surgery, University of Kansas School of Medicine-Wichita, Wichita, KS 67214, USA; Department of Biological Sciences, Wichita State University, Wichita, KS 67260, USA

**Keywords:** Osteosarcoma, DEPDC1/EEF1A1 complex, FOXO3a, regulatory pathway, poor prognosis

## Abstract

There are currently lack of effective therapeutic strategies for osteosarcoma, primarily due to insufficient understanding of the underlying mechanisms of the tumor cells. This study deciphers a potentially critical interplay of DEPDC1–EEF1A1–FOXO3a axis during the osteosarcoma progression. Bioinformatics analysis of documented 25,035 genes for differentially expressed genes were accompanied by transcriptional and translational examinations of clinical osteosarcoma specimens and osteosarcoma cell lines to assess the roles and interactions of DEP domain-containing 1 (DEPDC1), Elongation Factor 1-alpha 1 (EEF1A1), and FOXO3a in the tumor cells proliferation and prognosis. Gene expression profile analysis and clinical tests revealed highly expressed DEPDC1 in human osteosarcoma cells and tumor tissues. Vector-mediated silence of DEPDC1 resulted in halted osteosarcoma cell proliferation, promoted apoptosis, and ceased tumor metastasis. Immunoprecipitation assay confirmed that EEF1A1 directly bind to DEPDC1 protein through three binding regions. Further, DEPDC1/EEF1A1 complex significantly decreased the expression of FOXO3a at transcription and translation levels, which subsequently promoted the proliferation of osteosarcoma cells and tumor metastasis. Correlation studies exhibited that overexpression of DEPDC1/EEF1A1 complex in the clinical specimens negatively correlated with the patient survival rate. In conclusion, DEPDC1-EEF1A1–FOXO3a axis plays as a critical pathway that regulates the progression and prognosis of osteosarcoma.

## Introduction

Osteosarcoma is a common primary malignant bone tumor in children and adolescents. Surgical wide resection and drug chemotherapy are the main treatment for osteosarcoma. However, the side effects of chemotherapy and the drug resistance of osteosarcoma often lead to poor prognosis(Ritter J and Bielack SS 2010, Harrison DJ, Geller DS et al. 2018, Shen L, Zhao K et al. 2019). Thus, deciphering the molecular mechanisms underlying the osteosarcoma progression is of urgent.

Recent studies mainly focused on exploring the critical oncogenes that promote the occurrence and development of osteosarcoma. Because the occurrence and development of malignant tumor is a continuous and complex process, it involves a variety of changes in gene expression. A large number of studies have found that *XIAP*, *COX-2*, *Livin,* and other genes are highly expressed in osteosarcoma, while *Caspase-3* and *tp53* genes are low expressed in osteosarcoma(Huang FM, Chou MY et al. 2003, Notarbartolo M, Poma P et al. 2005, Spee B, Jonkers MD et al. 2006, Li X, Fan S et al. 2013, Wang L, Jin F et al. 2014). XIAP and Livin are main members of the apoptosis inhibitory protein family, can directly affect Caspase-3, Caspase-7i and Caspase-9 to result in inhibition of tumor cell apoptosis(Wunder JS, Gokgoz N et al. 2005, Spee B, Jonkers MD et al. 2006, Li X, Fan S et al. 2013, Wang L, Jin F et al. 2014). High expression of *COX-2* is mainly involved in promoting tumor angiogenesis and inhibiting tumor apoptosis(Huang FM, Chou MY et al. 2003). As a tumor suppressor gene, *tp53* gene induces apoptosis of tumor cells through Bax/Bcl2, Fas/Apol and IGF-BP3(Wunder JS, Gokgoz N et al. 2005). According to the change of spatial conformation, mutant p53 loses the regulatory effect on cell growth, apoptosis and DNA repair, and transforms from tumor suppressor gene to oncogene(Lee DF, Su J et al. 2015).

Therefore, it has been confirmed that the gene regulation network of osteosarcoma mainly suppresses the apoptosis of tumor cells, but the underlying mechanism in proliferation and metastasis of osteosarcoma is still unclear. The objectives of the current study tended to screen the differentially expressed genes that interact in promoting osteosarcoma cell proliferation and metastasis using strategies including bioinformatical analysis of NCBI Gene Bank data, assessment of clinical osteosarcoma specimens and an experimental osteosarcoma animal model, and *in vitro* mechanistic experimentations. Interestingly, a few genes are identified to be interactively associated with the progression of osteosarcoma.

DEP domain-containing 1 (DEPDC1) is a newly identified tumor-related gene. Recent studies have shown that DEPDC1 is overexpressed in bladder cancer, breast cancer, lung adenocarcinoma, colorectal cancer, and other malignant tumor types(Harada Y, Kanehira M et al. 2010, Kretschmer C, Sterner-Kock A et al. 2011, Okayama H, Kohno T et al. 2012, Obara W, Eto M et al. 2017, X. Shen and J. Han 2020). DEPDC1 inhibits cell apoptosis by activating the NF– κB pathway to promote the progression of bladder cancer(Harada Y, Kanehira M et al. 2010). However, there is no published evidence of whether DEPDC1 plays roles in the development and progression of human osteosarcoma. Preliminary data analysis of online database in our laboratory have suggested that DEPDC1 may also overexpressed in human osteosarcoma tissues. It appears worth to thoroughly investigate its roles and molecular pathways during the progression of osteosarcoma.

EEF1A1, a subtype of eukaryotic translation elongation factor 1A (EEF1A), is involved in the process of protein translation. Studies also identified that EEF1A1 can mediate the epithelial-mesenchymal transition (EMT) of breast cancer cells to promote the occurrence and metastasis through the formation of TGF-β-activated-translational (BAT) mRNP complex(Hussey GS, Chaudhury A et al. 2011). Slega E et al. found that EEF1A1 was overexpressed among the pancreatic cancer, leukemia and osteosarcoma cell lines, and siRNA targeting on EEF1A1 resulted in a chemo-sensitization toward Methotrexate (Selga E, Oleaga C et al. 2009). Recently, a bioinformatics analysis of osteosarcoma database indicated that EEF1A1 remains one of the 10 core genes in the interaction network during osteosarcoma tumorigenicity (Y. M. Ren, Y. H. Duan et al. 2018). However, it is still unclear how EEF1A1 is regulated to participate in the proliferation and metastasis of osteosarcoma.

FOXO3a belongs to the forkhead family of transcription factors that plays critical roles in regulating many cellular activities including cell cycle process, proliferation, and apoptosis. Loss of this critical transcription factor is associated with tumorigenesis and metastasis(Tenbaum SP, Ordóñez-Morán P et al. 2012, Levanon K, Sapoznik S et al. 2014, Liu Y, Ao X et al. 2018). Therefore, targeting on nuclear FOXO3a may develop potential therapeutic means for certain types of cancers(M. Tarrado-Castellarnau, R. Cortes et al. 2015, Yao S, Fan LY et al. 2018) However, it is elusive how FOXO3a functions in osteosarcoma.

In this study, we revealed that DEPDC1 combined with EEF1A1 to diminish the expression of FOXO3a. The DEPDC1-EEF1A1 complex also promotes phosphorylation of FOXO3a, ultimately accelerates osteosarcoma cell proliferation and migration *in vitro*. Understanding and characterization of the DEPDC1–EEF1A1–FOXO3a axis may provide a meaningful and effective target for treating human osteosarcoma.

## Results

### DEPDC1 is highly expressed in human osteosarcoma tissues and cells

To explore the crucial oncogenes related to osteosarcoma progression, we adopted bioinformatics techniques to analyse the differential gene expressions between an osteosarcoma group and a normal group from the Gene Expression Omnibus (GEO) database including 25,035 annotated genes. A total of 1,355 differentially expressed genes were screened out (523 upregulated genes and 832 downregulated genes) **(Fig. 1A**). For the combined adjusted *P*- and logFC values, the logFC value of *DEPDC1* was the largest (logFC = 1.204), with a corresponding adjusted *P*-value <0.05. It is suggested that the relative expression level of *DEPDC1* in the osteosarcoma group was 2.3 times higher than that in the control group (**Fig. 1B**). Next, the protein and RNA expression levels of DEPDC1 in the human osteosarcoma specimens and the osteosarcoma cell lines (HOS, MG-63, and Saos-2) were compared with the adjacent normal tissues and the human osteoblastic cell line (Hfob1.19), indicating a significantly higher expression levels at both post-transcriptional and post-translational levels in the tumor tissue and osteosarcoma cells than those in normal controls (**Fig. 1C–E**).

**Fig. 1.**
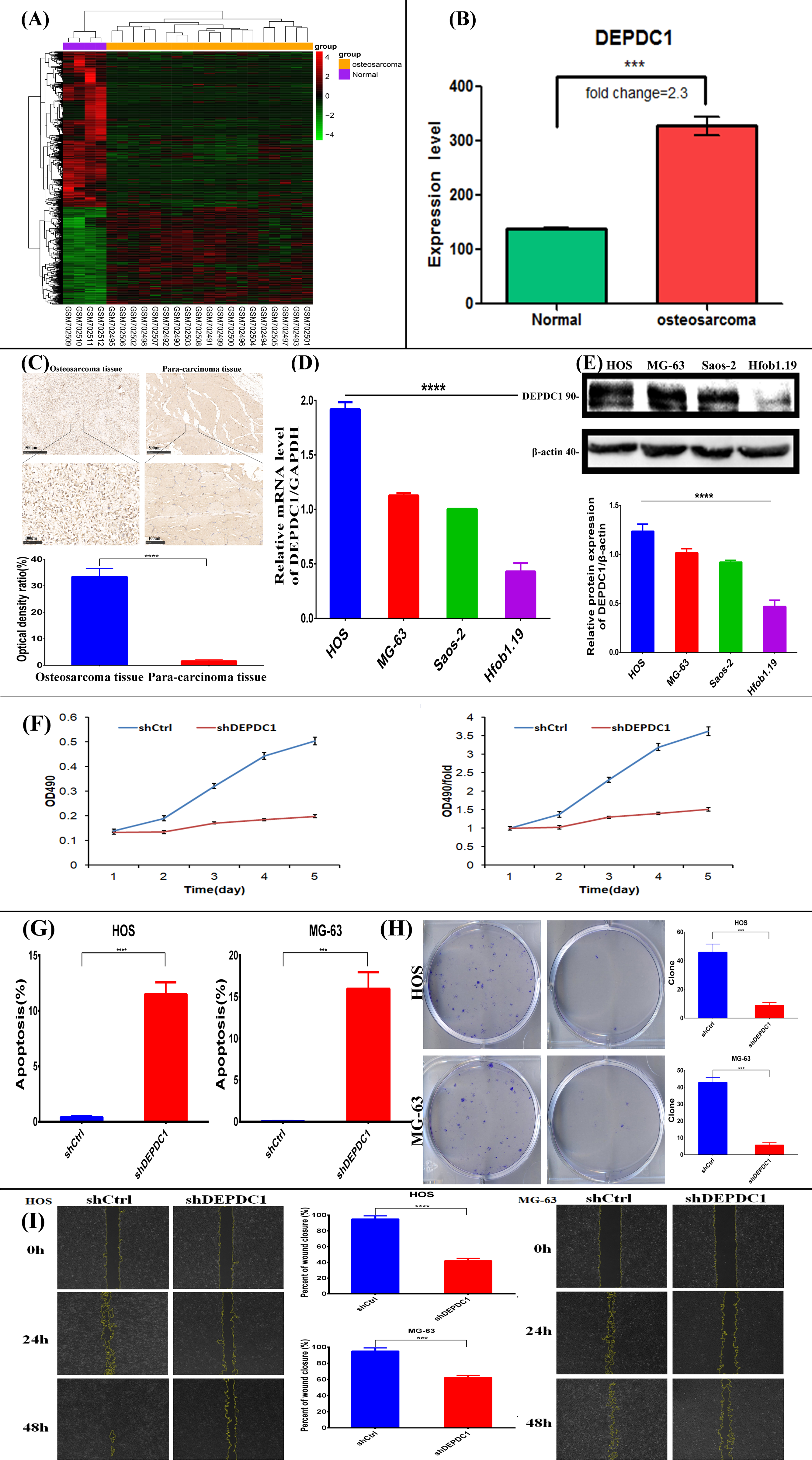
DEPDC1 is highly expressed in GEO database and human osteosarcoma tissues and cells, meanwhile, down-regulation of DEPDC1 significantly inhibits the growth, metastasis and promoted the apoptosis of osteosarcoma cells. **A, B** Heat map and bar chart showing the expression levels of DEPDC1 across the human osteosarcoma among GEO database. C Representative immunohistochemical (IHC) staining of DEPDC1 protein expression classified in osteosarcoma patient tissues. Scale bar, 100 μm. **D, E** RT-PCR and Western blot analysis of DEPDC1 expression in osteosarcoma cells and normal human osteoblasts. **F** MTT assay showed that knockdown of DEPDC1 expression in HOS cells significantly suppressed the cell proliferation. **G** Flow cytometry was used to determine the apoptosis percentages of each group. **H** Cell clone formation of HOS and MG-63 cell lines after knockdown of DEPDC1. **I** The cell scratch assay was used to examine the metastatic abilities of HOS and MG-63 osteosarcoma cells after inhibiting the expression of DEPDC1. ***p<0.001,****p<0.001.

### Suppression of DEPDC1 expression significantly inhibits the proliferation and metastasis, but promote apoptosis of osteosarcoma cells

Lentiviral vectors-mediated silence of *DEPDC1* expression in osteosarcoma cells resulted in the diminished cell growth (cell counts by MTT assay, **Fig. 1F**), increased apoptotic cell ratio (Annexin V Staining, **Fig. 1G**), repressed cell colonies formation (clonogenicity assay, **Fig. 1H**), and halted cell migration (scratch assay, **Fig. 1I**). Taken together, these findings demonstrate that DEPDC1 is an oncogene to promote the osteosarcoma cancer cell growth and cell migration while inhibit cell apoptosis.

### DEPDC1 promotes the proliferation and migration of osteosarcoma cells by binding to and upregulating EEF1A1 expression

To determine the specific signaling pathway of DEPDC1 in the proliferation and metastasis of osteosarcoma cells, immunoprecipitation (IP) and mass spectrometry were performed to fish out the candidate proteins that directly interact with DEPDC1. It appears that EEF1A1 is one of the most likely proteins to bind DEPDC1 **(Table. 1**) (Z Gu, J Xia et al. 2017, FP McManus, F Lamoliatte et al. 2017). Co-immunoprecipitation (Co-IP) experiments showed that EEF1A1 and DEPDC1 formed a complex in osteosarcoma cells (**Fig. 2A, 2B)**. To identify the binding site of DEPDC1 to EEF1A1, we constructed four plasmids encoding for different *DEPDC1* fragments (15–106, 107–180, 181–406, and 407–527) (**Supplementary Table S1**). We co-transfected FLAG-tagged-*DEPDC1* and His-tagged-*EEF1A1* into 293T cells and HOS cells. DEPDC1 and its four fragments were successfully expressed after transfection, as detected with a FLAG antibody (**Fig. 2C top**). The fragments of 15–106, 107–180, and 407–527 showed strong binding with His-tagged-*EEF1A1*, while the binding was lost in 181–406 (**Fig. 2C bottom**). These results suggested that DEPDC1 binds to EEF1A1 at three domains, 15–106, 107–180, and 407–527. Subsequently, we silenced EEF1A1 expression with siRNAs targeting the open reading frame of *EEF1A1* (siEEF1A1; **Supplementary Table S1**). Silencing of *EEF1A1* can reduce the proliferation (**Fig. 2D**) and migration (**Fig. 2E**) induced by overexpression of DEPDC1 in both HOS and MG-63 cells. Collectively, these results demonstrated that DEPDC1 interacts with EEF1A1, which is crucial for DEPDC1 promotion on the proliferation and migration of osteosarcoma cells.

**Table. 1.**
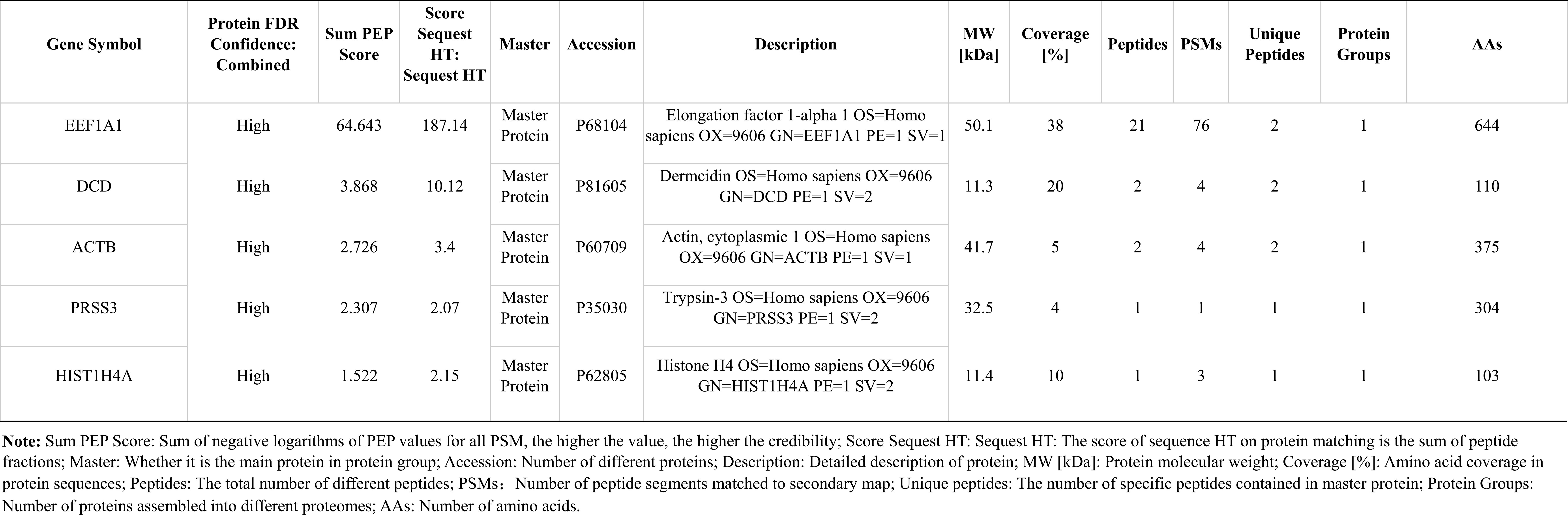
Protein identification list of samples.

**Fig. 2.**
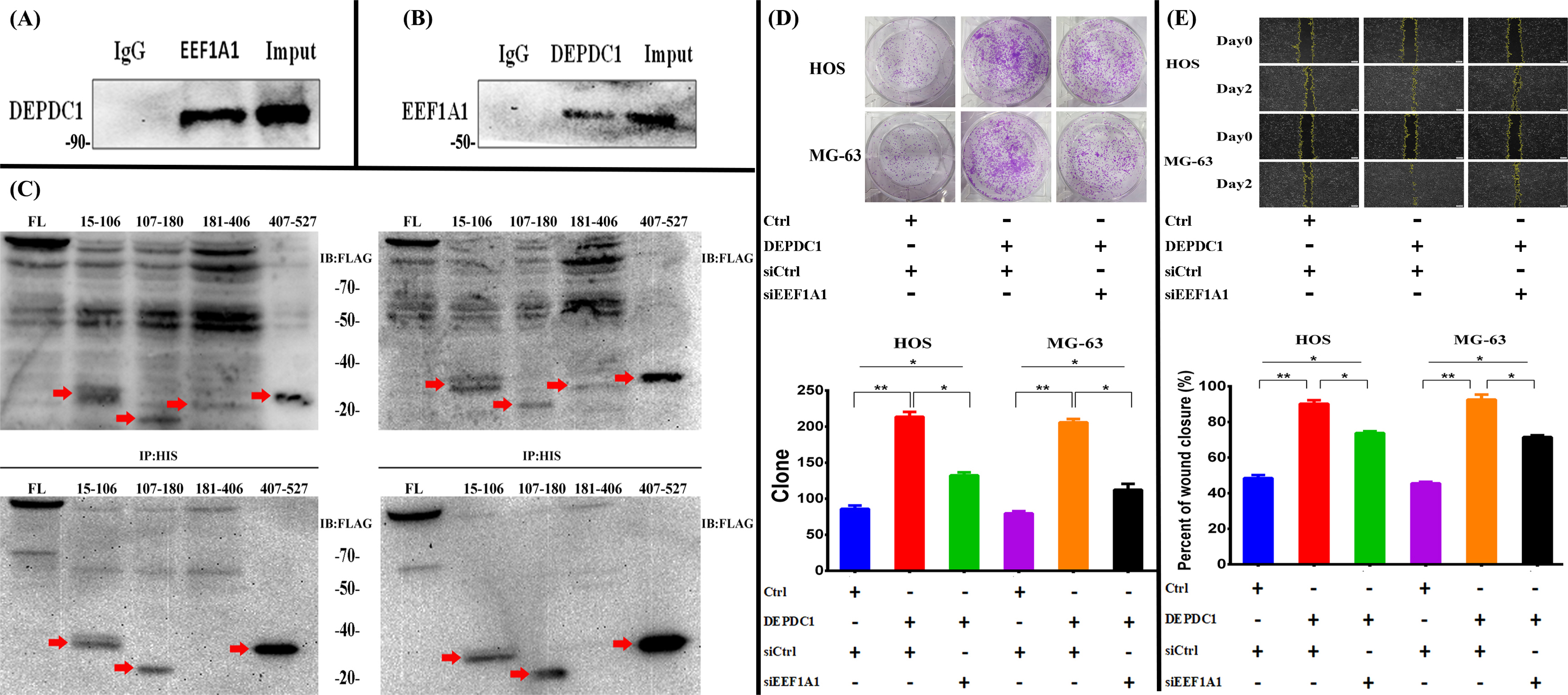
DEPDC1 enhances the growth and metastasis of osteosarcoma cells by binding to the extension factor 1-alpha 1 (EEF1A1) and promoting the expression of EEF1A1. Co-immunoprecipitation (Co-IP) of DEPDC1 **(A)** and EEF1A1 **(B)** in osteosarcoma cells. **C** Co-IP assay to detect the binding site of DEPDC1 to EEF1A1 in 293T cells and HOS cells. Plasmids of FLAG-tagged DEPDC1 and its N-terminal deletion mutants or C-terminal deletion mutants and His-tagged EEF1A1 were co-transfected. Cell lysates were immunoprecipitated with His antibody and probed with FLAG antibody. Expression of FLAG-tagged N-terminal deletion or C-terminal deletion mutants of DEPDC1 probed by FLAG antibody after transfection for 48h (top). Co-IP assay immunoprecipitated (bottom) with His antibody and probed with FLAG antibody. The binding regions of DEPDC1 and EEF1A1 were 15-106,107-180 and 407-527. **D, E** Cell clone formation and cell scratch assay were used to examine the proliferation and metastatic abilities of HOS and MG-63 osteosarcoma cells, while knockdown of EEF1A1 after DEPDC1 overexpression.*p<0.05, **p<0.01.

### DEPDC1 inhibits FOXO3a expression by regulating the binding of EEF1A1 to DEPDC1

To further explore the signalling pathway involved in the promotion of DEPDC1–EEF1A1 complex on human osteosarcoma, we performed RNA-seq to investigate the differentially expressed genes in *DEPDC1*-knockdown cells (**Supplementary Fig. S1E**). Transcription factor enrichment analysis with TFactS showed that FOXO3 as a transcription factor was significantly regulated by DEPDC1 (**Supplementary Fig. S1H**). Lenti-viral vector-mediated overexpression of DEPDC1 on osteosarcoma cells (HOS and MG-63 cells) resulted in a significantly decrease of FOXO3a expression, but an elevated cyclin D1 expression. However, knockdown the expression of DEPDC1 obtained reverse responses of FOXO3a and cyclin D1 expressions at both mRNA and protein levels (**Fig. 3A, B**). Furthermore, after downregulating DEPDC1, the resulted low expression of FOXO3a accompanied with the diminished apoptotic activities of osteosarcoma cells and active cell migration (**Fig. 3C, D**).

**Fig. 3.**
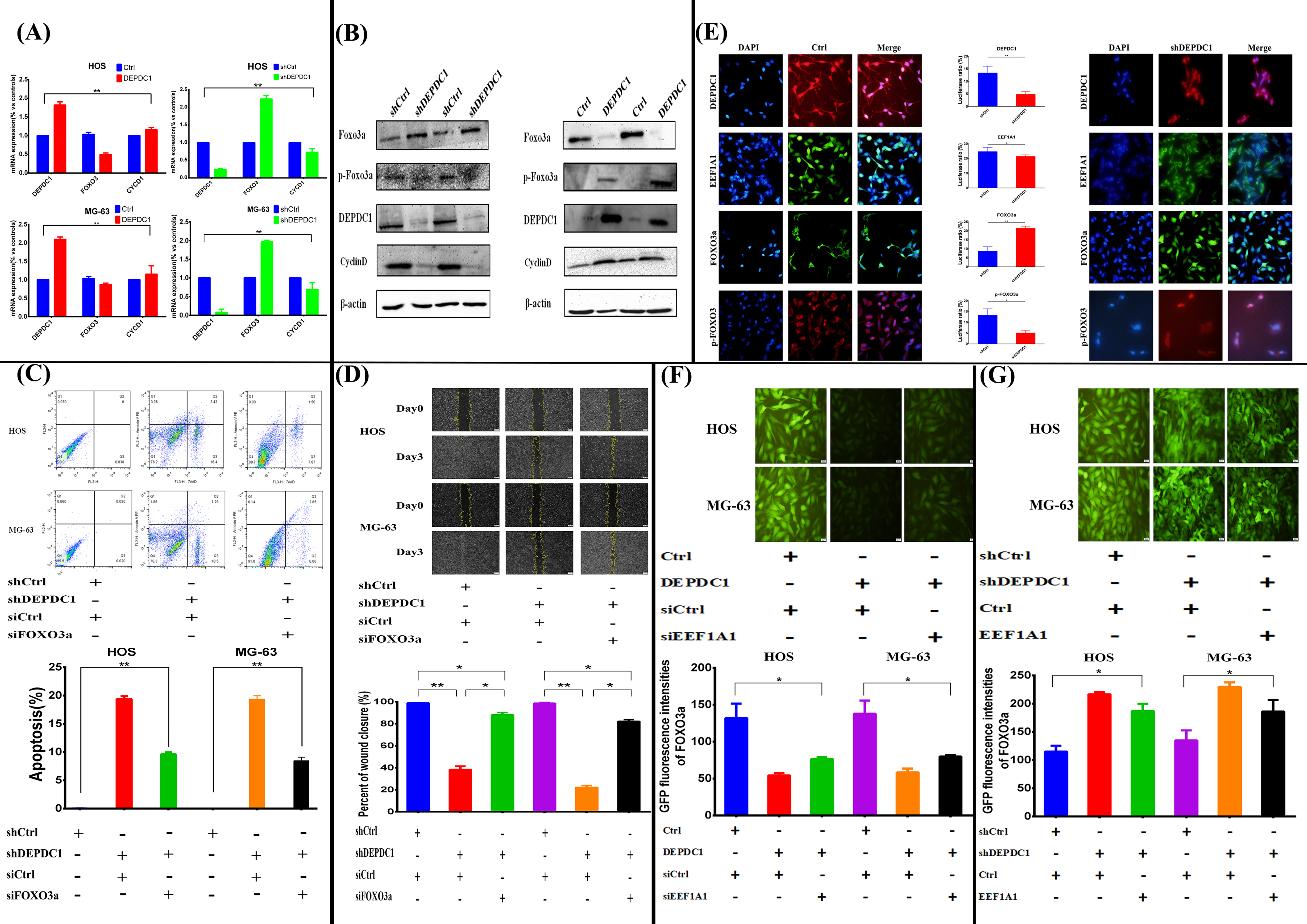
Down-regulation of DEPDC1 inhibits the metastasis and promotes the apoptosis of osteosarcoma cells by up-regulating forkhead box O3 activation (FOXO3a). **A, B** Real-time PCR and Western blot analysis were used to analyze the correlation between DEPDC1 and FOXO3a pathway after up-regulation and down-regulation of DEPDC1 gene in HOS and MG-63 cells. After down-regulating DEPDC1, FOXO3a was down-regulated in HOS and MG-63 osteosarcoma cells for flow cytometry of apoptosis **(C)** and cell scratch assay **(D)**. **E** Immunofluorescence staining of DEPDC1, EEF1A1, FOXO3a and p-FOXO3 in Ctrl group and shDEPDC1 group. **F, G** The GFP fluorescence intensity of FOXO GFP Reporter Plasmid regulated by DEPDC1 and EEF1A1. *p<0.05, **p<0.01.

Histology analysis indicated that DEPDC1 and EEF1A1 were co-localized in the nucleus to support the complex formation of DEPDC1 and EEF1A1 (**Fig. 3E left**), while FOXO3a and p-FOXO3a were largely colocalized in the cytoplasm. However, the cytoplasm staining of FOXO3a was dramatically diminished and returned to the nuclei of osteosarcoma cells after knockdown of DEPDC1 expression (**Fig. 3E right**). Quantification of the Luciferases activity suggested that FOXO3a was up-regulated in the cytoplasm of the cells with silence of DEPDC1 expression, while EEF1A1 and p-FOXO3a appeared diminished compared to wild-type (Controls) cells, in the nucleus and cytoplasm, respectively (**Fig. 3E middle**). To further explore the regulatory effect of DEPDC1 and EEF1A1 on FOXO3a, a FOXO3-GFP reporter plasmid (**Supplementary Fig. S1I**) was used to transfect to HOS and MG-63 cells to illustrate FOXO3 levels. Compared with the control group, overexpression of DEPDC1 dramatically diminished the FOXO3-GFP fluorescence intensity. However, blockade of EFF1A1 on the high DEPDC1 cells can partially recover the GFP fluorescence intensity of FOXO3 (**Fig. 3F**). Conversely, when DEPDC1 was downregulated, the GFP fluorescence intensity of FOXO3 was significantly enhanced, and the GFP fluorescence intensity of FOXO3 was decreased while increasing EEF1A1 expression (**Fig. 3G**). These results suggest that the combination of DEPDC1 and EEF1A1 promotes FOXO3a to translocate from the nucleus for phosphorylation, thus promoting the migration of osteosarcoma cells and inhibiting their apoptosis.

### Perturbed expression of DEPDC1 inhibits the proliferation of osteosarcoma cells in vivo

To investigate the role of DEPDC1 in the progression of osteosarcoma in vivo, a nude mouse xenograft model was established with infusion of wild-type or DEPDC stably knockdown (shDEPDC1 transduced) osteosarcoma cells. The experimental osteosarcoma tumor sizes from the DEPDC1-knockdown human osteosarcoma cells were significantly smaller than those from wild type osteosarcoma cells (**Fig. 4A, C, D**). Hematoxylin and eosin (H&E) staining showed that the *DEPDC1*-knockdown tumors presented with multi-areas of necrosis, cell nuclear condensations and cytoplasmic fibrosis (**Fig. 4B**). IHC staining confirmed that both DEPDC1 and EEF1A1 expressions were diminished. whereas expression of FOXO3a was markedly elevated, in the *DEPDC1*-knockdown tumors (**Fig. 4E, F**).

**Fig. 4.**
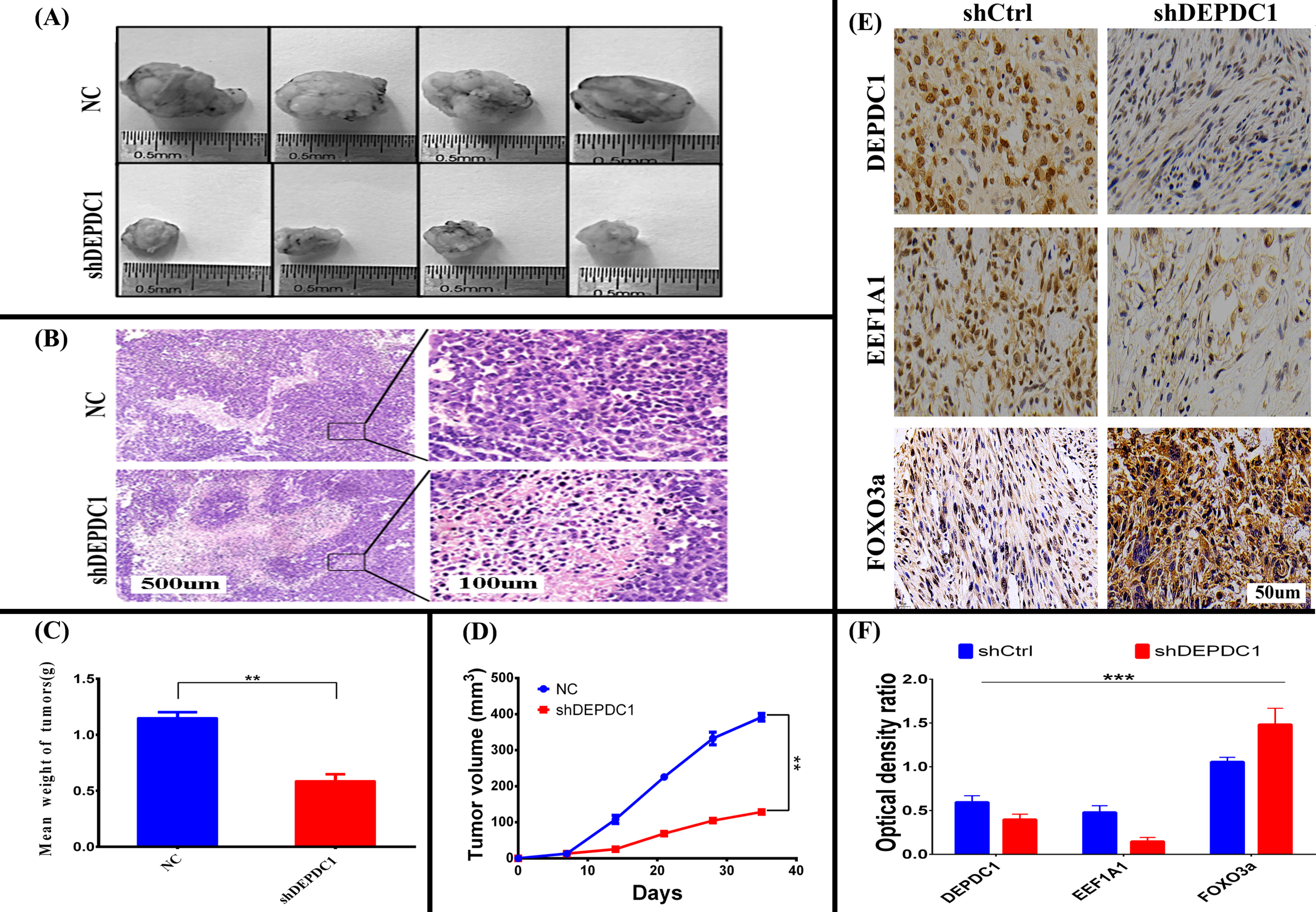
Down-regulation of DEPDC1 inhibits the growth of osteosarcoma cells in vivo. **A** Representative primary tumor volumes and tumor images are shown. **B** Representative H&E staining images of the two groups. Scale bar, 100 μm. Mean tumor weight **(C)** and tumor volume **(D)** of NC group and shDEPDC1group. **E** Representative IHC image of xenograft tumors for DEPDC1, EEF1A1 and FOXO3a in shCtrl group and shDEPDC1group. **F** Quantitative results were derived from **(E)**, using Image J software. **p<0.01, ***p<0.001.

### DEPDC1, EEF1A1, and FOXO3a expression levels correlate with human osteosarcoma progression

To explore the relationship between the DEPDC1/EEF1A1/FOXO3a axis and the clinicopathological characteristics of osteosarcoma patients, we analysed the positive staining patterns of DEPDC1, EEF1A1, and FOXO3a in clinical human osteosarcoma tissue specimens by immunohistochemistry (**Fig. 5A**). The data clearly shown that the expression of DEPDC1 protein was positively correlated with EEF1A1 but negatively correlated with FOXO3a, which indicated that the development of human osteosarcoma may be related to the DEPDC1/EEF1A1– FOXO3a pathway (**Fig. 5B**). Further, the IHC data was correlated with the clinical disease progressions of the patients. The protein expression levels of either DEPDC1 or EEF1A1 were higher in the advanced TNM stage group and the lymphatic metastasis-positive group. However, FOXO3a was expressed at lower levels in either the advanced TNM stage or lymphatic metastasis-positive groups (**Fig. 5C**). Subsequently, receiver operating characteristic (ROC) curves demonstrated that the area under the curves (AUC) of the DEPDC1-, EEF1A1-, and FOXO3a-based predictions were 0.7908, 0.79, and 0.7528, respectively, suggesting that they all could be used to predict the survival rate of osteosarcoma patients (**Fig. 5D**). Importantly, the higher expression levels of either DEPDC1 or EEF1A1 and lower expression of FOXO3a were highly associated with the decreased survival time of osteosarcoma patients (*P*<0.001, **Fig. 5E**). Indeed, the survival time was shortest in the high expression groups of DEPDC1 and EEF1A1 (**Fig. 5F**) and the groups with high DEPDC1 but low FOXO3a expression (**Fig. 5G**). **Fig. 5H** indicated that the shortest survival time were among the patients with high expression of both DEPDC1 and EEF1A1 but low expression of FOXO3a. Furthermore, the χ^2^ value of log-rank (Mantel-Cox) test in the three-index group is 74.05, demonstrating that combination of the three indices would be a better predictor for progression of osteosarcoma patients.

**Fig. 5.**
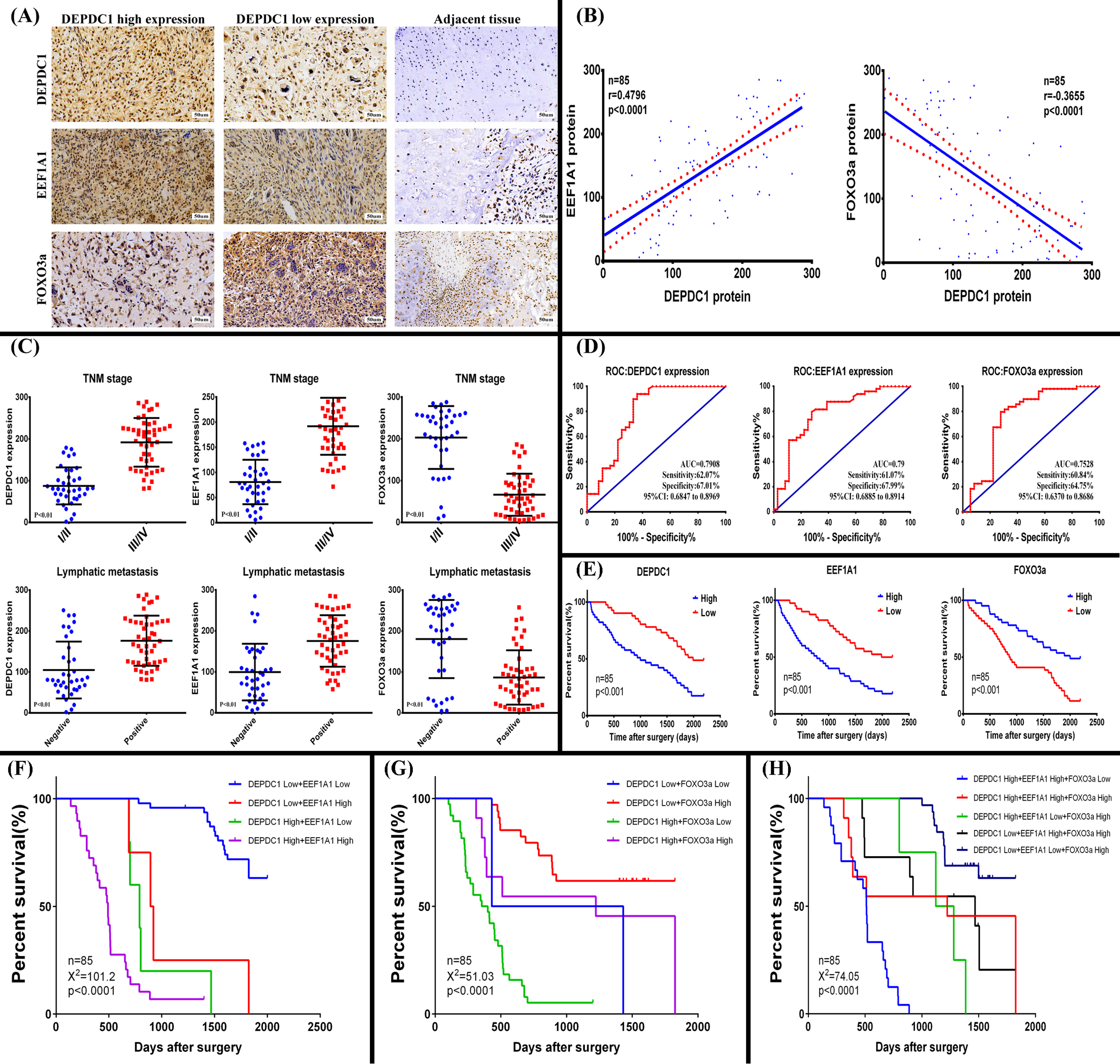
The expression of DEPDC1 in human osteosarcoma is consistent with EEF1A1, but negatively correlated with FOXO3a. **A** Representative immunohistochemical staining image of DEPDC1, EEF1A1, FOXO3a in osteosarcoma and related adjacent non-cancerous tissues. **B** The correlation between the protein levels of DEPDC1 and EEF1A1 (left). The correlations between the protein levels of DEPDC1 and FOXO3a (right) (p<0.0001). **C** The association between DEPDC1/ EEF1A1/ FOXO3a expression and TNM stage/ lymphatic metastasis in patients with osteosarcoma (p<0.01). **D** The receiver operating characteristic (ROC) curves for predicting patients’ survival time using DEPDC1/ EEF1A1/ FOXO3a expression. **E** Kaplan–Meier analyses of overall survival according to DEPDC1/ EEF1A1/ FOXO3a expression levels (p<0.01). AUC, area under the curve; TNM, tumor node metastases. **F, G** Kaplan-Meier analyses of overall survival time combining two indices (p<0.0001). **H** Kaplan-Meier analyses of overall survival time combining three indices (p<0.0001).

## Discussion

In this study, we explored a new mechanism of DEPDC1-induced proliferation and migration of osteosarcoma cells. The expression of DEPDC1 in osteosarcoma was similar to that of EEF1A1 but was negatively correlated with FOXO3a. Therefore, we speculated that the complex formed by DEPDC1 and EEF1A1 enhanced the expression of EEF1A1 and ultimately led to the downregulation of FOXO3a, thus promoting the proliferation and migration of osteosarcoma cells in vitro and in vivo (**Fig. 6**).

**Fig. 6.**
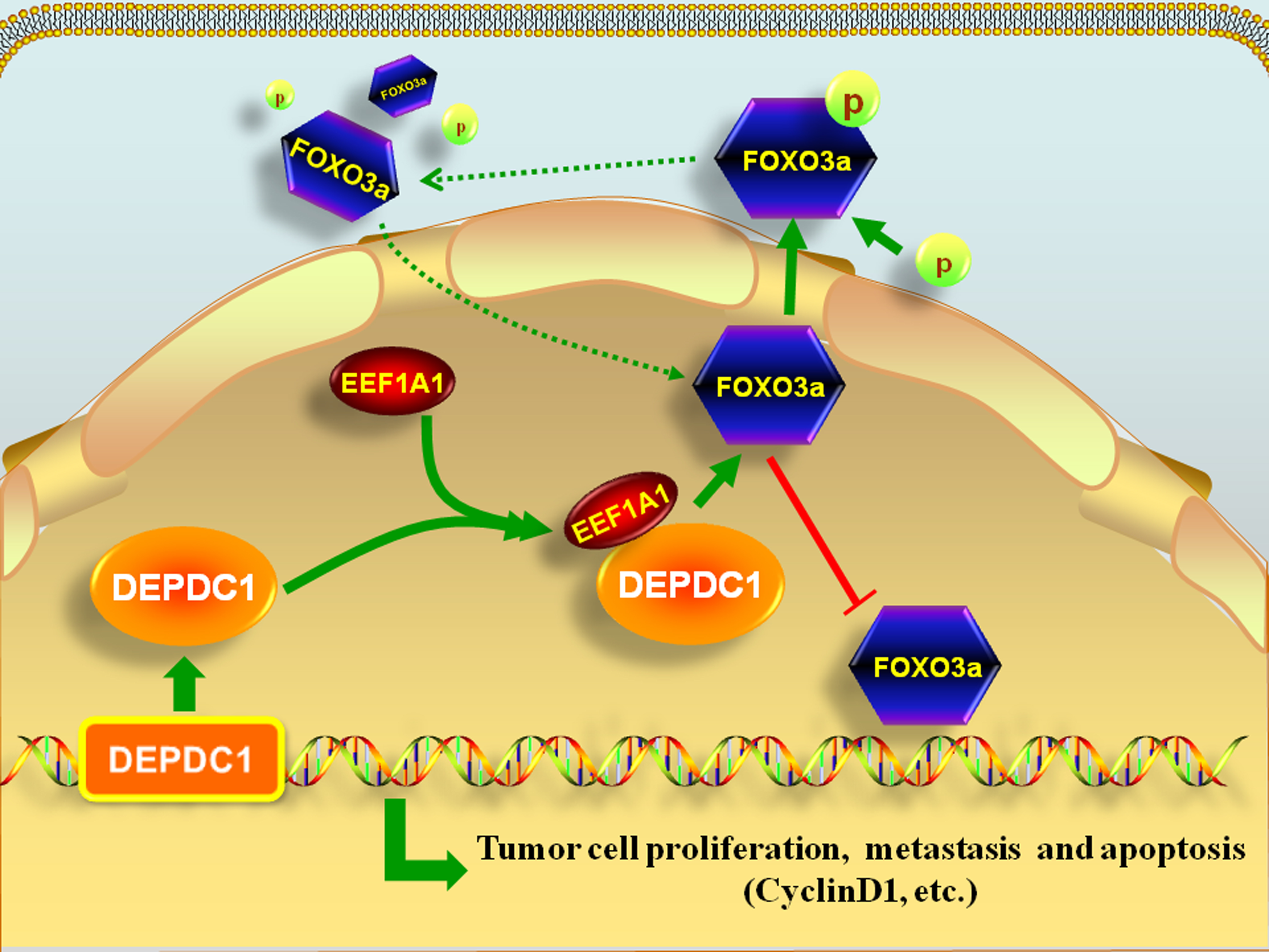
A proposed model of the signaling pathway for DEPDC1 regulating the human osteosarcoma proliferation, metastasis and apoptosis. In this model, DEPDC1 improves the expression of EEF1A1 and enhances the binding to EEF1A1, thus form a complex to inhibit the FOXO3a which could block tumor progression. This process ultimately promotes the proliferation, migration and inhibits the apoptosis of osteosarcoma.

*DEPDC1* is a newly discovered tumor-related gene that has a highly conserved domain. Many studies have found that proteins with DEP domains can regulate many cellular functions, such as cell membrane anchoring, signal transduction, cell polarity establishment, and regulation of small molecule GTP enzyme activity(Spring K, Chabot C et al. 2012). Recent studies have shown that DEPDC1 is overexpressed in bladder cancer, breast cancer, lung adenocarcinoma, and other malignant tumor types(Harada Y, Kanehira M et al. 2010, Kretschmer C, Sterner-Kock A et al. 2011, Okayama H, Kohno T et al. 2012). In addition, Harada et al. found that DEPDC1 mainly inhibited cell apoptosis through the NF–κB signalling pathway, and then promoted the progression of bladder cancer Cell-permeable peptide DEPDC1-ZNF224 interferes with transcriptional repression and oncogenicity in bladder cancer(Harada Y, Kanehira M et al. 2010). Furthermore, DEPDC1 can also be used in the diagnosis and treatment of various tumors. Kretschmer et al. found that DEPDC1 and FOXM1 are significantly upregulated in ductal carcinoma in situ (DCIS), and thus can be used to identify early molecular markers of breast cancer(Zhang L, Du Y et al. 2019). S-288310, a cancer peptide vaccine containing oncoantigens against DEPDC1, is well tolerated and can effectively prolong the survival time of patients with urothelial carcinoma of the bladder (Obara W, Eto M et al. 2017). However, it remains unclear whether *DEPDC1* is the main mechanism of promoting the proliferation and migration of malignant tumors. In this study, we found that *DEPDC1* was highly expressed in human osteosarcoma through the GEO database analysis and confirmed this result in osteosarcoma tissues and cells **(Fig. 1**). At the same time, through co-IP and RNA-seq experiments, we found that *DEPDC1* inhibits the expression of FOXO3a in combination with EEF1A1, thereby promoting the proliferation and migration of osteosarcoma in vitro and in vivo **(Fig. 2–5**).

Eukaryotic translation elongation factor 1A (EEF1A) is an important molecule involved in the translation function in protein synthesis. EEF1A can be divided into two subtypes, EEF1A1 and EEF1A2. In humans, EEF1A is encoded by two genes on chromosomes 6 and 20, which are mainly involved in apoptosis, cell cycle regulation, protein degradation, and post-translation modifications(Kapp LD and Lorsch JR 2004, Andersen CB, Becker T et al. 2006, Behrmann E, Loerke J et al. 2015). EEF1 complex members are necessary in the eukaryotic elongation process.

Recently, it has been found that EEF1A1 is related to cancer occurrence(Selga E, Oleaga C et al. 2009, Liu S, Hausmann S et al. 2019). Genetic changes in *EEF1A1* were detected by The Cancer Genome Atlas (TCGA) to explore its potential impact on selected epigenetic modulators. However, the specific upstream and downstream regulatory molecules that bind to EEF1A1 were not accurately investigated, nor the clinical correlation between the expression of EEF1A1 and the prognosis of tumor patients(Tayou J, Wang Q et al. 2016). Furthermore, the role of EEF1A1 in the proliferation and metastasis of osteosarcoma has not been studied. In our study, we found that DEPDC1 directly binds to and promotes the expression of EEF1A1 in the nuclei of osteosarcoma cells, thus promoting the proliferation and migration of osteosarcoma cells in vitro and in vivo (**Fig. 2 and 4**).The 15–106, 107–180, and 407–527 fragments of *DEPDC1* showed strong binding with EEF1A1 (**Fig. 2C bottom**). Furthermore the 15–106 domain of DEPDC1 is the DEP domain, named after Dishevelled, Egl-10, and Pleckstrin, in which this domain was first discovered. The function of this domain remains unclear, but is believed to be important for the membrane association of the signalling proteins in which it is present(Lishko PV, Martemyanov KA et al. 2002, Jonsson H and Peng SL 2005). In addition, this report for the first time revealed the relationship between the expression of DPEDC1/EEF1A1 and the clinical prognosis of osteosarcoma patients (**Fig. 5**). The expression of EEF1A1 in osteosarcoma is positively correlated with DEPDC1, and the high expression of both can reduce the survival time of osteosarcoma patients, which indicates that DEPDC1 and EEF1A1 are potential prognostic markers and therapeutic molecular targets of osteosarcoma.

The Forkhead transcription factors (FOXO), also named Forkhead-like protein (FKHR), is a family transcription factors that was identified in 2000. There are four types in mammals, FOXO1, FOXO3a, FOXO4, and FOXO6, which are distributed on different chromosomes(2017).

The common feature of this family is the conserved DNA domain, namely, the Forkhead box. This protein family regulates apoptosis, cell cycle, cell proliferation, DNA damage repair, and cancer development, and inhibits tumor cell proliferation(Tayou J, Wang Q et al. 2016). FOXO3a is among the most widely studied members of the Forkhead family. It is located on human chromosome 6q21 and is expressed in gastrointestinal, liver, ovary, prostate, and breast tissue as well as others(Tran H, Brunet A et al. 2002, Mao Z, Liu L et al. 2007). FOXO3a dysfunction leads to uncontrolled cell proliferation and DNA damage accumulation, resulting in tumorigenic effects(Hu C, Ni Z et al. 2017, Liu Y, Ao X et al. 2018). The main mechanism regulating FOXO3a activity and its target genes is the control of the nuclear–cytoplasmic shuttling of FOXO3a. The phosphorylation of FOXO3a leads to its translocation from the nucleus to cytoplasm, followed by binding to 14-3-3 protein in the cytoplasm, and then FOXO3a is degraded in a ubiquitin-/proteasome-dependent manner(Tsai KL, Sun YJ et al. 2007, Kapoor I, Li Y et al. 2019). The balance of nuclear–cytoplasmic shuttling is crucial for maintaining the function of FOXO3a. The loss of this balance leads to the occurrence and development of various diseases including cancer(Levanon K, Sapoznik S et al. 2014, Fluckiger A, Dumont A et al. 2016). Studies have shown that the Wnt–β-catenin and PI3K–AKT–FOXO3a pathways have a central role in cancer. AKT phosphorylates FOXO3a, promoting its translocation from the cell nucleus to the cytoplasm. When this effect was reversed by PI3K and AKT inhibitors, the accumulation of FOXO3a in the nucleus increased, which promoted the apoptosis of colon cancer cells and inhibited their metastasis (Hu MC, Lee DF et al. 2004). Additionally, Hu et al. found that nuclear exclusion of FOXO3a by AKT contributed to cell survival. They also observed that Iκβ kinase (IKK) can promote FOXO3a phosphorylation, inhibiting the expression of FOXO3a, and finally causing FOXO3a protein hydrolysis through the ubiquitin-dependent proteasome pathway. The expression and accumulation of FOXO3a in the nucleus is reduced by IKKβ, which promotes the proliferation of breast cancer cells and is related to the low survival rate of breast cancer(George J, Lim JS et al. 2015). However, there have been no further studies on whether oncogenes are involved in the transcription and expression of FOXO3a, affecting the balance of nuclear–cytoplasmic shuttling and promoting the occurrence and development of tumors. Lei et al. found that *DEPDC1* overexpression facilitated cell proliferation and tumor growth through increasing the expression of FOXM1 in TNBC cells (Zhang L, Du Y et al. 2019). FOXM1 is negatively regulated by FOXO3a, supports cell survival, drug resistance, colony formation and proliferation in vitro, and promotes tumor development in vivo(Buchner M, Park E et al. 2015). At the same time, the FOXO3-FOXM1 axis is a key cancer drug target and a modulator of cancer drug resistance(Yao S, Fan LY et al. 2018). But so far, there is no report to explore the relationship between *DEPDC1* and FOXO3a and their interaction mechanism. In this study, we found that *FOXO3* had the best correlation with *DEPDC1* expression in human osteosarcoma cells by RNA-seq (**Fig. 3A, B**). Later, we found that DEPDC1 forms a complex with EEF1A1. Upregulation of DEPDC1 promoted the accumulation of EEF1A1 in the nuclei of osteosarcoma cells and then inhibited the expression of FOXO3a, restricting its distribution to the cytoplasm, while the expression of phosphorylated FOXO3a increased (**Fig. 2–4**). When DEPDC1 was downregulated, the opposite result was obtained (**Fig. 2–4**). Furthermore, the expression of FOXO3a was negatively correlated with DEPDC1 in human osteosarcoma and lowexpression of FOXO3a shortened the survival time of patients with osteosarcoma (**Fig. 5**). Therefore, we observed that inhibition of FOXO3a expression could significantly promote tumor proliferation and migration while reducing the survival time of tumor patients, which is consistent with many previous reports. Wasim et al. found that EEF1A1 can activate AKT dependent cell migration and tumor proliferation(Lin KW, Yakymovych I et al. 2010, Abbas W, Kumar A et al. 2015). In addition, AKT phosphorylates FOXO3a, promoting tumor proliferation and metastasis(Giamas G, Filipović A et al. 2011).

Cyclin D1 is an important regulator of the cell cycle and has a vital role in tumor development. In the cell cycle, cyclin D1 levels are regulated by many factors and change periodically. Cyclin D1 regulates cell proliferation from G1 phase to S phase. Overexpression of cyclin D1 shortens the time from G1 to S phase and accelerates the transformation of the cell cycle, leading to uncontrolled cell proliferation and migration and finally tumor occurrence(Tudzarova S, Trotter MW et al. 2010). In addition, cyclin D1 is the critical downstream molecule of the FOXO3a signalling pathway and is negatively correlated with FOXO3a expression (**Fig. 3A, B**). Tudzarova et al. found that FOXO3a activates the ARF–Hdm2–p53–p21 pathway, and p53 in turn activates expression of the Wnt/β-catenin signalling antagonist DKK3, leading to cyclin D1 downregulation(Tudzarova S, Trotter MW et al. 2010). Zheng et al. also found that FOXO transcription factors repress cyclinD1 transcription. Failure to hydroxylate FOXO3a promotes its accumulation in cells, which in turn suppresses cyclin D1expression(Zheng X, Zhai B et al. 2014). Lin et al. found that *FLOT1* knockout inhibited the proliferation and tumorigenicity of breast cancer cells by upregulating FOXO3a and then downregulating cyclin D1(Lin C, Wu Z et al. 2011). Meanwhile, the overexpression of EEF1A1 regulates G1-phase progression to promote HCC proliferation through the STAT1-cyclin D1 pathway(Huang J, Zheng C et al. 2017). STAT1 has been proved to have the same effect as AKT, which can phosphorylate FOXO3a(Sanjay de Mel, Susan Swee-Shan Hue et al. 2019). However, it has not been reported whether the complex formed by DEPDC1 and EEF1A1 in human osteosarcoma weakens the inhibitory effects of FOXO3a on cyclin D1 to ultimately affect tumor progression.

Combined with this study and many previous studies, we have confirmed that the proliferation and migration of osteosarcoma cells are affected through the DEPDC1–EEF1A1–FOXO3a– cyclin D1 signalling pathway in vitro and in vivo. Therefore, we postulate that during development of human osteosarcoma, high expression of DEPDC1 permits it to form a complex with EEF1A1 and promote the expression of EEF1A1, thus inhibiting the synthesis of FOXO3a. At the same time, FOXO3a is translocated from the nucleus and gets phosphorylated, which eventually decreases the inhibitory effect of FOXO3a on cyclin D1 and promotes the growth of human osteosarcoma (**Fig. 6**). However, there are some limitations in this study, such as how FOXO3a is degraded after leaving the nucleus and how it affects the expression of downstream cyclin D1 in osteosarcoma, thus inhibiting the progression of osteosarcoma. Further studies are warranted to elucidate the mechanisms.

## Conclusion

In this study, we deciphered the mechanism of DEPDC1 promoting the development of osteosarcoma, which will provide new therapeutic targets for further development of new anticancer drugs. At the same time, DEPDC1–EEF1A1–FOXO3a axis can accurately predict the clinical characteristics and prognosis of patients, thus providing new methods and strategies for tumor diagnosis and treatment.

## Materials and Methods

### Gene Expression Omnibus data analysis of differential gene expression in human osteosarcoma

Eight datasets (species: *Homo sapiens*) were downloaded from the NCBI Gene Expression Omnibus (GEO) database including GSE11414, GSE12865, GSE14359, GSE16088, GSE19276, GSE28424, GSE36001, and GSE9508. The Limma software package (v. 3.10.3) was used to analyse the differences in gene expression between the osteosarcoma group and normal group to obtain the corresponding *P*- and logFC values. The Benjamini–Hochberg method(Yoav Benjamini and Yosef Hochberg 1995) was performed for multiple tests and corrections, and the corrected *P*-value (namely the adjusted *P*-value) was obtained.

### Clinical samples and survival time analysis

Total of 85 Osteosarcoma tissue specimens including their adjacent normal tissues were obtained, along with the de-identifiable clinical chart information, from Jinan Central Hospital, Qilu Hospital of Shandong University, and Xi’an Best Biotechnology Co., Ltd. This multi-institutional collaborative human investigation project was approved by the individual institutional ethical committees. Receiver operating characteristic (ROC) curve was used to analyse the relationship between protein expression and survival time. The values of ROC curves with the maximized sum of sensitivity and specificity were used as the thresholds to separate the high and low expression of DEPDC1, EEF1A1, and FOXO3a, and the survival time of each group was compared.

### Cell culture and reagents

Human osteosarcoma cell lines HOS, MG-63, and Saos-2 were obtained from GENECHEM Co., Ltd (Shanghai, China), and the human osteoblast cell line Hfob1.19 was obtained from Zhong Qiao Xin Zhou Biotechnology Co., Ltd (Shanghai, China). All cell lines were cultured in Dulbecco’s modified Eagle’s medium supplemented with 10% foetal bovine serum, streptomycin (100 mg/mL), and penicillin (100 U/mL).

### Lentiviral and plamid vectors constructs and transfection

GV-based lentiviral vectors coding for DEPDC1 (Lenti-DEPDC1) and empty vectors (Lenti-Ctrl) were designed and constructed by GENECHEM Co., Ltd (Shanghai, China). DEPDC1 siRNA and shRNA plasmids, as well as DEPDC1 shRNA Lentiviral particles (shPEDC1) were also obtained from the same vendor for DEPDC1 silence studies. The lenti-DEPDC1, lenti-Ctrl, shDEPDC1 and shCtrl viral or plasmid vectors were co-cultured with the osteosarcoma cells at a multiplicity of infection (MOI) of 25, 25, 20, and 20 respectively. The core sequences of target gene fragments in each group are shown in a Supplementary (Table S1).

In addition, four segments of DEPDC1 full-length sequence (15–106, 107–180, 181–406, 407–527) were incorporated into plasmids for EEF1A1 binding assays. They were constructed in FLAG-tagged plasmids, while *EEF1A1* plasmids were HIS-tagged. The plasmids containing siEEF1A1, siFOXO3a, and FOXO3-GFP reporter plasmid were also constructed (Table S1).

### Immunoprecipitation

293T and HOS cells were co-transfected with FLAG-tagged-*DEPDC1*, its four segments (15– 106, 107–180, 181–406, 407–527), and HIS-tagged-*EEF1A1* plasmids for 24 h. The cells were lysed in Radio-Immunoprecipitation Assay (RIPA) Lysis Buffer (P0013K, Beyotime Biotechnology, Shanghai, China), 1 mg of extracted protein was transferred to Eppendorf tubes to incubate with anti-FLAG-antibody (1:1000; Millipore Sigma) or anti-HIS-antibody (1:300; Santa Cruz Biotechnology) for immunoprecipitation.

### Immunofluorescent (IF) staining

Cells were cultured to 60%–80% density on glass coverslips in culture dishes. After discarding the culture medium, the cells were washed three times with PBS, fixed with 4% paraformaldehyde at 25°C for 20min, and permeabilized with 0.2% Triton X-100 at 25°C for 10min. After washing with PBS, cells were incubated with anti-DEPDC1-antibody (1:200; Novus Biologicals), anti-EEF1A1-antibody (1:1000; Millipore Sigma), anti-FOXO3a-antibody (1:200; Cell Signaling Technology), or anti-Phospho-FOXO3a-antibody (1:200; Invitrogen Antibodies), respectively, at 8°C overnight. The cells were next washed with PBS three times and incubated with a fluorescent dye-conjugated secondary antibody (1:300; Thermo Fisher Scientific) and phalloidin at 37°C for 2 h. After rinse in PBS three times, fluorescent staining of the cells was observed using a Zeiss LSM710 confocal microscope (Carl Zeiss AG, Oberkochen, Germany).

### Cell proliferation and clonogenic survival assays

Cells infected with either shCtrl or DEPDC1-siRNA lentivirus were cultured at 5x10^4^/well in 96-well plate at 37°C and 5% CO_2_ for 5 days before tested for MTT cell proliferation assay as described previously(S. Yang, K. Zhang et al. 2015). For the clonogenic assay, 1×10^3^ HOS and MG-63 cells infected with shDEPDC1 or shCtrl lentivirus (or DEPDC1, Ctrl, and DEPDC1+siEEF1A1) were inoculated into each well of six-well culture plates. The cells were incubated at 37°C and 5% CO_2_ for 14 days. The cell colonies were fixed with 4% paraformaldehyde and stained with Giemsa. Colonies consisting of *>*50 cells were counted.

### Apoptosis analysis

HOS or MG-63 cells were seeded into six-well plates at a density of 1.5×10^5^cells/well, incubated for 24 h, and divided into shCtrl, shDEPDC1, and shDEPDC1+siFOXO3a groups. After 24 h, the cells from all groups were harvested and washed twice with cold PBS. Apoptotic cells were distinguished by Annexin V-7AAD/PI dual staining and an apoptosis detection kit from Keygen Biotech (Nanjing, China). Flow cytometry was also performed to detect apoptotic cells (FCM, BD FACSCalibur; BD Biosciences). All experiments were performed in triplicate.

### Wound healing assay

A total of1.5×10^6^ HOS or MG-63 cells in log phase were seeded into each well of a six-well plate and divided into shCtrl, shDEPDC1, shDEPDC1+ siFOXO3a, DEPDC1, Ctrl, and DEPDC1+ siEEF1A1 groups. When the cell density reached approximately 80%, cells from all four groups were rinsed twice with phosphate-buffered saline (PBS) and wounded (scratched) with a pipette tip. The cells were imaged and the total initial wounded area, S0h, was calculated using ImageJ software (NIH, MS, USA). Cells were cultured in serum-free medium for 48h. The cells were imaged and the total final wounded area, S48h, and the percentage of wound closure (1-[S0h–S48h]*/* S0h) were calculated.

### Protein detection by mass spectrometry

Q Exactive TM LC-MS system from Thermo Co. Ltd. was used for mass spectrometry analysis. The peptide samples were aspirated by an automatic injector and combined with a C18 capture column (3μm, 75μm × 20mm, 100Å), and then eluted to the analysis column (50μm × 150mm, 2μm particle size, 100Å Pore size, Thermo) for separation. Two mobile phases (mobile phase A: 1% DMSO, 99% H2O, 0.1% formic acid and mobile phase B: 1% DMSO, 80% ACN, 0.1% formic acid) were used to establish a 100 minute analytical gradient (0 min in 3% B, 7 min of 3-5%; 65 min of 5-18% B, 10 min of 18-33% B, 2 min of 33-90% B, 90% B for 6 min). The flow rate of the liquid phase was set at 300 nL / min. In DDA mode analysis of mass spectrometry, each scan cycle contains one MS full scan (m / z range is 350-1800, ion accumulation time is 200 ms), followed by 40 MS / MS scans (m / z range is 100-1500, ion accumulation time is 50 ms). The condition of MS / MS acquisition is that the parent ion signal is greater than 3e6 and the charge number is + 2 ∼ + 5. The exclusion time of repeated ion collection was set to 35 s.

The mass spectrum data generated by QE was retrieved by Protein Discover (V2.2), and Percolator was used as the database retrieval algorithm. The database used in the search is the human proteome reference database in UniProt (UniProt_Human_20180405.fasta). The retrieval parameters are as follows: Scan Event: Msaa Analyzer (FTMS), MS Order(MS2), Activation Type (HCD), scan typr(full); Sequest HT: Enzyme (Trypsin), Dynamic Modification (Oxdation, Acetyl, Carbamidomethyl). The Maximum Delta Cn and Maximum Rank of PSM ≥ 0.05 were selected as the criteria. Deleted the entries retrieved from the database and eliminated the information of contaminated proteins. Then the remaining identification information was used for subsequent analysis.

### RNA sequencing

Total RNA was extracted using Trizol reagent (Invitrogen). Total RNA was quantified by NanoDrop ND-2000 (Thermo Scientific) and detected using an Agilent Bioanalyzer 2100 (Agilent Technologies). Feature Extraction software (version 12.0.3.1, Agilent Technologies) was used to process the original images and extract the original data. Then we used Genespring software (version 14.8, Agilent Technologies) to standardize the data. After the standardized data were filtered, at least one group of 100% labelled probes detected in each group of samples for comparison were left for subsequent analysis. The *P*-values of t-tests were used to screen the differential genes, and the standard of screening was *P*≤0.05. Then, GO and KEGG enrichment analyses were performed to determine the biological functions or pathways that were mainly affected by the differentially expressed genes. Finally, the expression patterns of differential genes in different samples were displayed in the form of a thermogram(Y Gao, S Shang et al. 2020).

### TF analysis of transcription factors (TFs) regulated by downregulation of DEPDC1 in osteosarcoma

The microarray data of DEPDC1 downregulation in osteosarcoma were uploaded to the TFactS database at (http://www.tfacts.org)(A Essaghir, F Toffalini et al. 2010). The TFs were predicted using the P-value, q-value, E-value and false discovery rate (http://www.tfacts.org/); only TFs that were <0.05 for all 4 indexes were considered to be reliable. The number of TFs mediated by down regulated DEPDC1 was counted. Subsequently, the number of shared TFs and different TFs of target genes were calculated and compared.

### Animal xenograft tumorigenesis assay

All animal experimentation protocols were approved by the Ethics Committee of Shandong University. Four- week-old female BALB/C nude mice (Beijing Vital River Laboratory Animal Technology Co. Ltd., Beijing, China) were randomly injected in the right subaxillary region either with a suspension of 6×107 wild-type MG-63 cells (12 mice, NC group) or shDEPDC1-infected MG-63 cells (12 mice, shDEPDC1 group). Body weight and tumor diameter were measured every other day. Tumor volume was calculated according to the formula (length [L] × width [W]2)/2. At day 37, the tumors were harvested, weighed, and fixed in 10% formalin before processed for paraffin-embedded tissue sectioning. The 5-μm-thick sections were stained for haematoxylin and eosin (H&E) and IHC. No tumor volume was >2000 mm^3^.

### In vitro assay of interplay of DEPDC1-EEF1a1-FOXO3 axis

The FOXO3-GFP reporter plasmid was co-transfected with siEEF1A1 plasmid into either the stable DEPDC1-overexpressing osteosarcoma cell lines or the DEPDC1-knockdown stable cell lines for 48 hours. GFP fluorescence intensity of the cells was observed using a Zeiss LSM710 confocal microscope (Carl Zeiss AG, Oberkochen, Germany) and analysed by ImageJ software (v1.47, NIH, MS, USA).

### Statistical analysis

Data were processed in GraphPad Prism 6.0 (GraphPad Software, CA, USA) and SPSS v. 20.0 (IBM Corp., NY, USA). The Kaplan–Meier method was used for survival analysis. For comparisons of two groups, Student’s t-test was applied if no significantly different variances were present; otherwise, the Mann–Whitney test was used. One-way ANOVA with Bonferroni’s test was used to compare >2 group means. Data are expressed as means ± SD. Differences were considered statistically significant at *P*<0.05.

## Acknowledgements

The authors would like to thank the Department of Orthopedic Surgery and Central Lab of Jinan Central Hospital for assistance.

## Abbreviations

DEPDC1: DEP domain-containing 1
EEF1A1: Eukaryotic translation extension factor 1A1
EMT: Epithelial-mesenchymal transition
FOXO: Forkhead transcription factor
GEO: Gene Expression Omnibus
TCGA: The Cancer Genome Atlas

## Author contributions

L Shen and H Li performed the experiments, analyzed the data and initiated the manuscript. C Zhou, Y Zhang, K Zhao, M Bretches, and L Lu participated in the experiments and manuscript preparation. H Li contributed to the animal model. A Zhang, R Liu, SY Yang and B Ning conceived the study and participated in manuscript preparation.

## Availability of data and materials

The datasets used and/or analyzed during the current study are available from the corresponding authors per request.

## Ethics approval and informative consent

All the clinical studies were approved by the Institutional Ethical Review Boards of Jinan Central Hospital, and written informed consent was obtained from all the patients or their families. All animal experiments were carried out in accordance with the guidelines approved by the Institutional Animal Care and Use Ethics Committee of Shandong University (SYXK20150015).

## Consent for publication

All authors have agreed to publish this manuscript.

---

**Supplementary Fig. S1.**
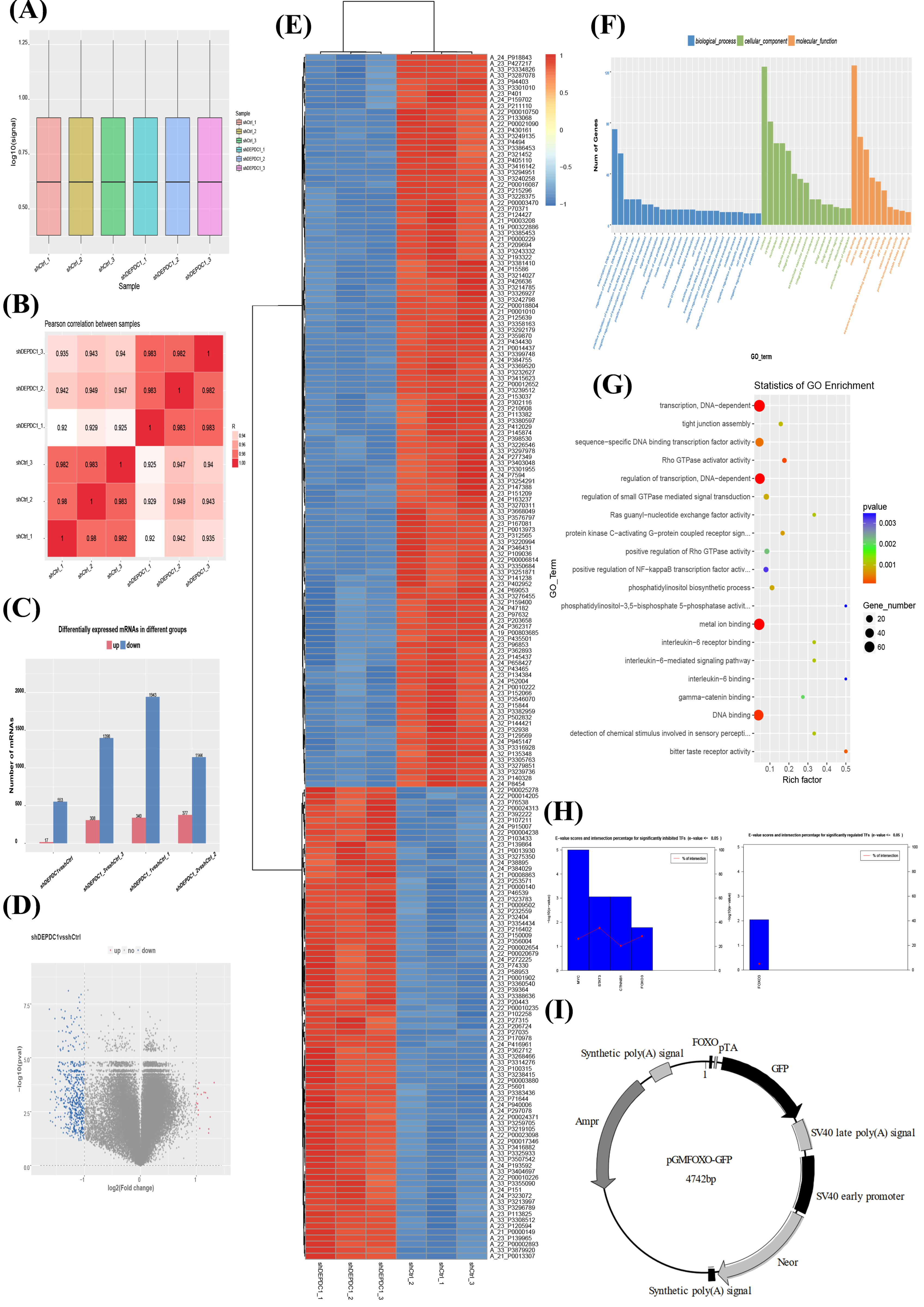
FOXO3a plays a major role in regulating the downstream signaling pathway of DEPDC1. **A, B** Boxplot of RNA expression and Person RNA correlation between samples in DEPDC1-knockdown HOS cells were revealed by Agilent gene expression profile chip analysis. The significant differences in up-regulated and down-regulated expressions in the differential expression analysis of each group were statistically analyzed and displayed in a bar chart (**C**). The overall distribution of differentially expressed genes can be known from the volcano map (**D**). In the analysis of differential expression mRNA clustering, top 200 with the minimum P value was selected as the heat map (**E**). Red is high expression gene, blue is low expression gene. Histogram (**F**) and scatter plot (**G**) of GO enrichment after inhibiting the expression of DEPDC1. **H** E−value scores and intersection percentage for significantly regulated TFs showed that FOXO3a was the best candidate molecule for the downstream regulation of DEPDC1. **I** Vector map of FOXO-GFP reporter plasmid.

**Supplementary Table S1.**
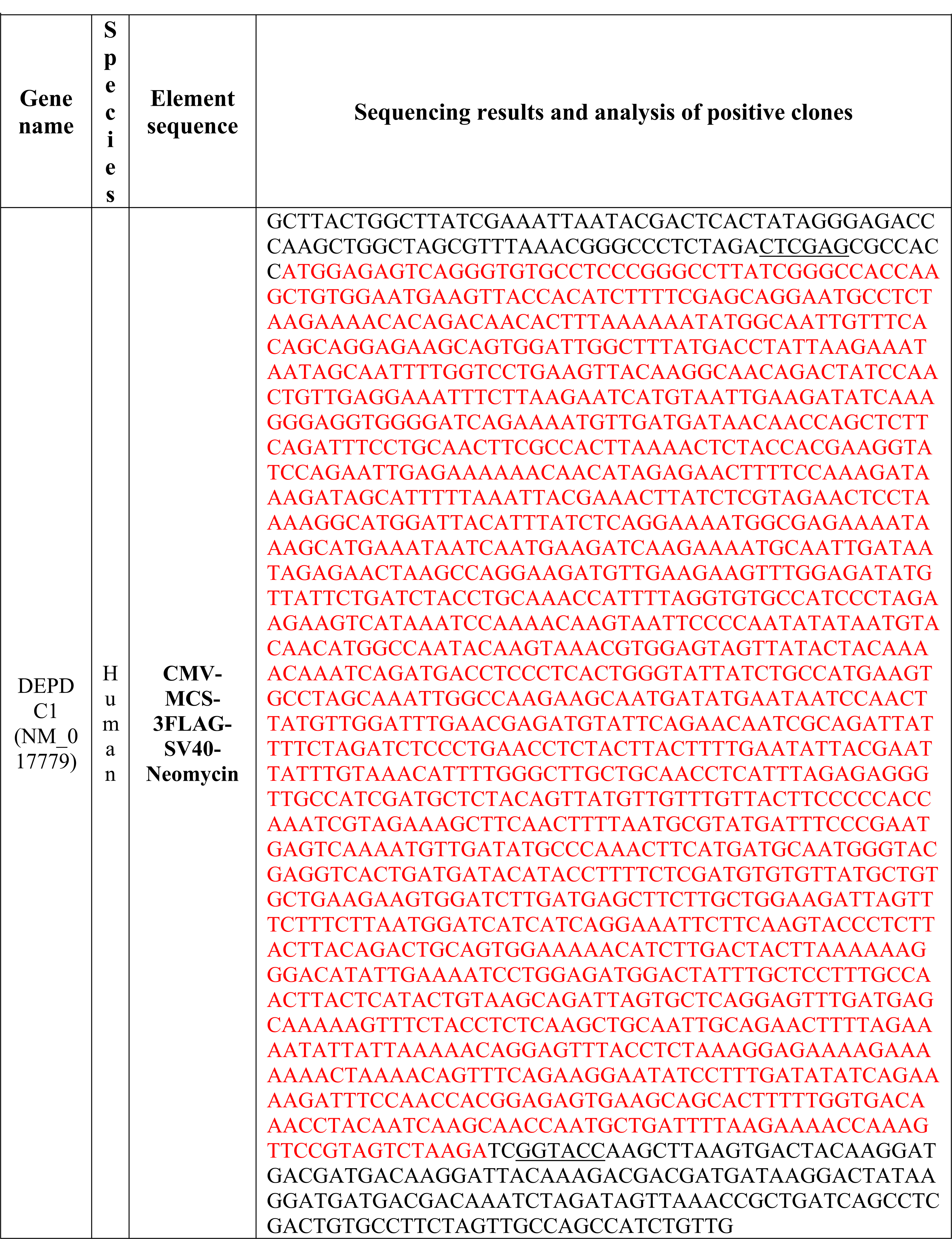
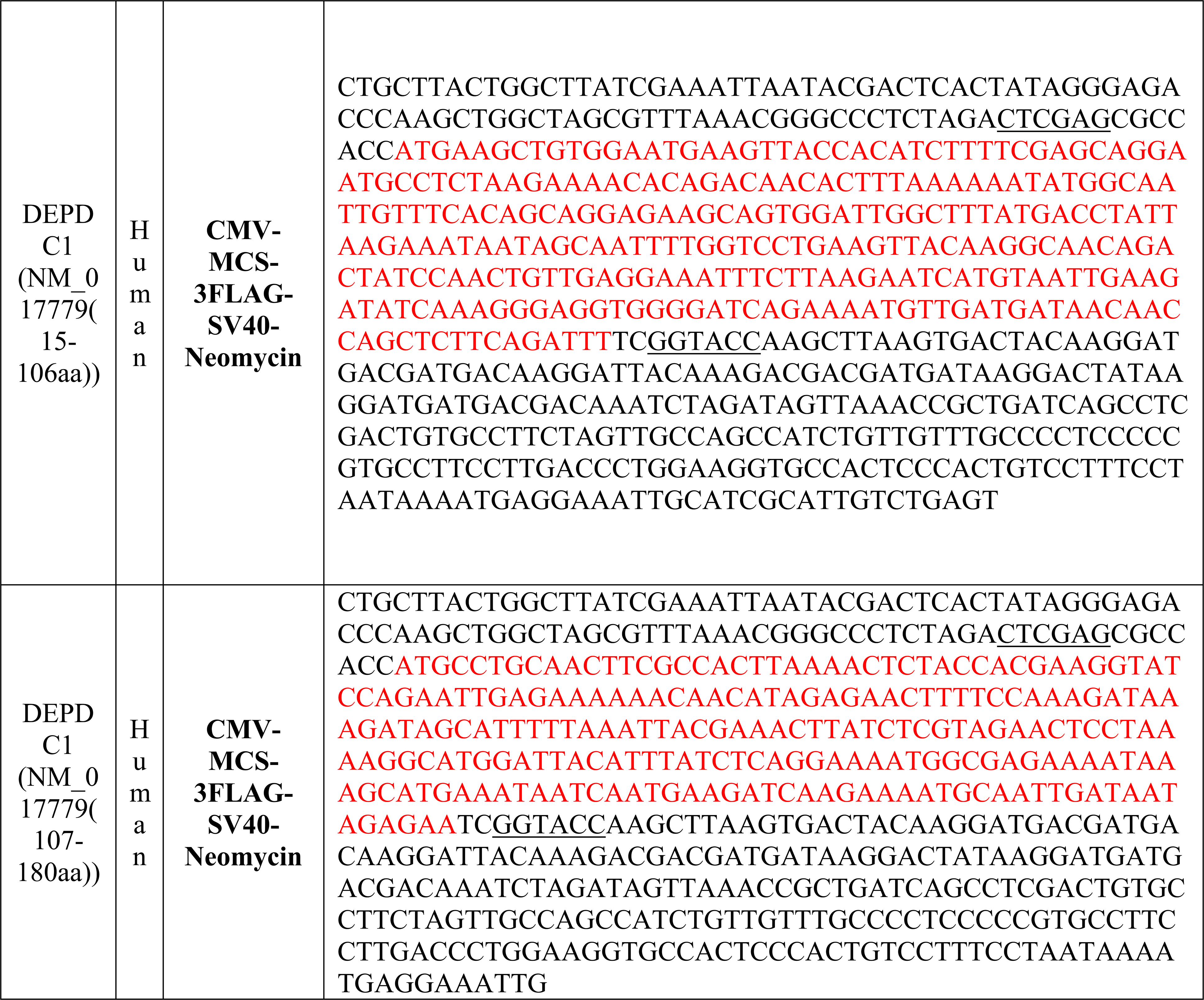
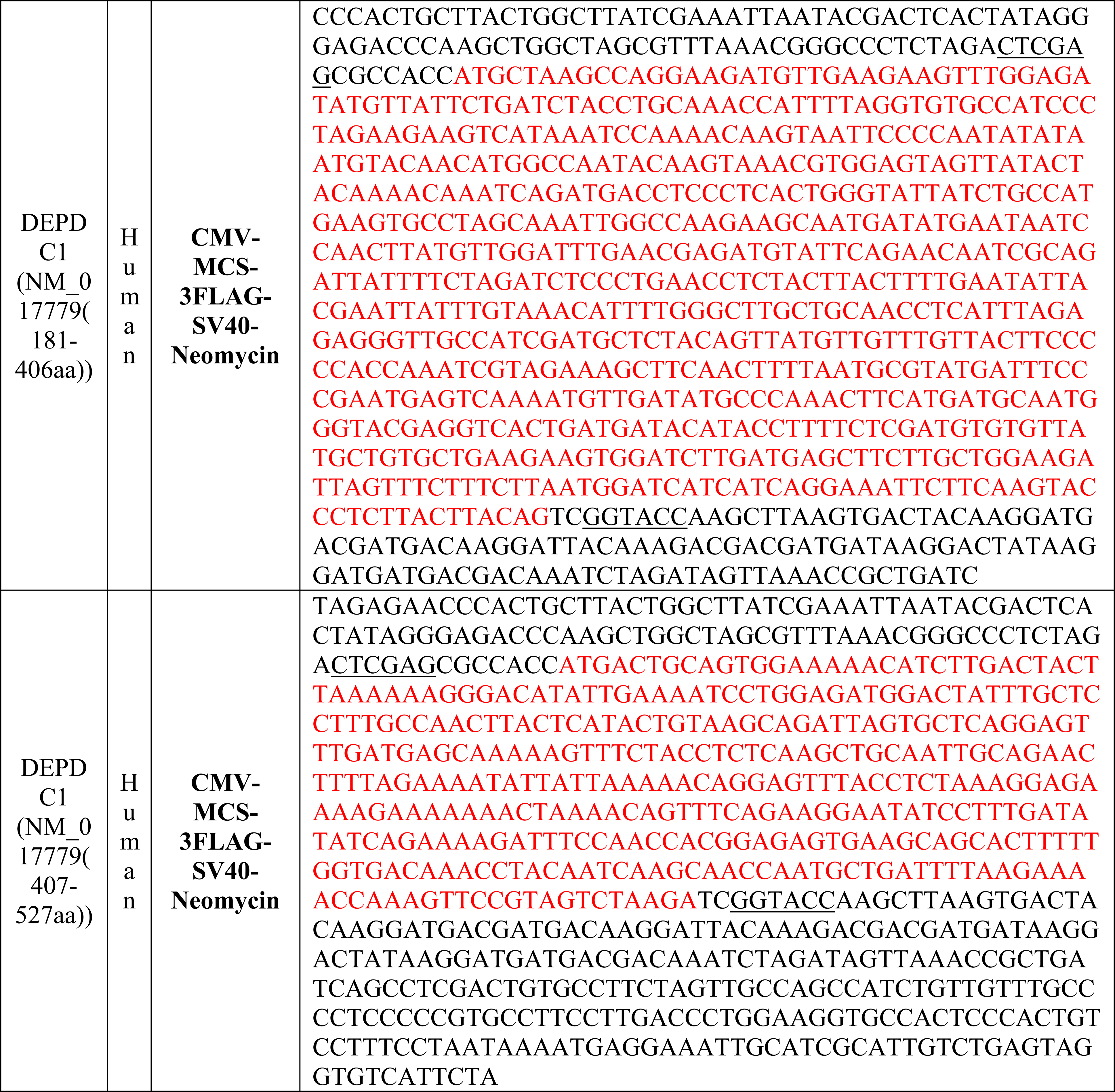
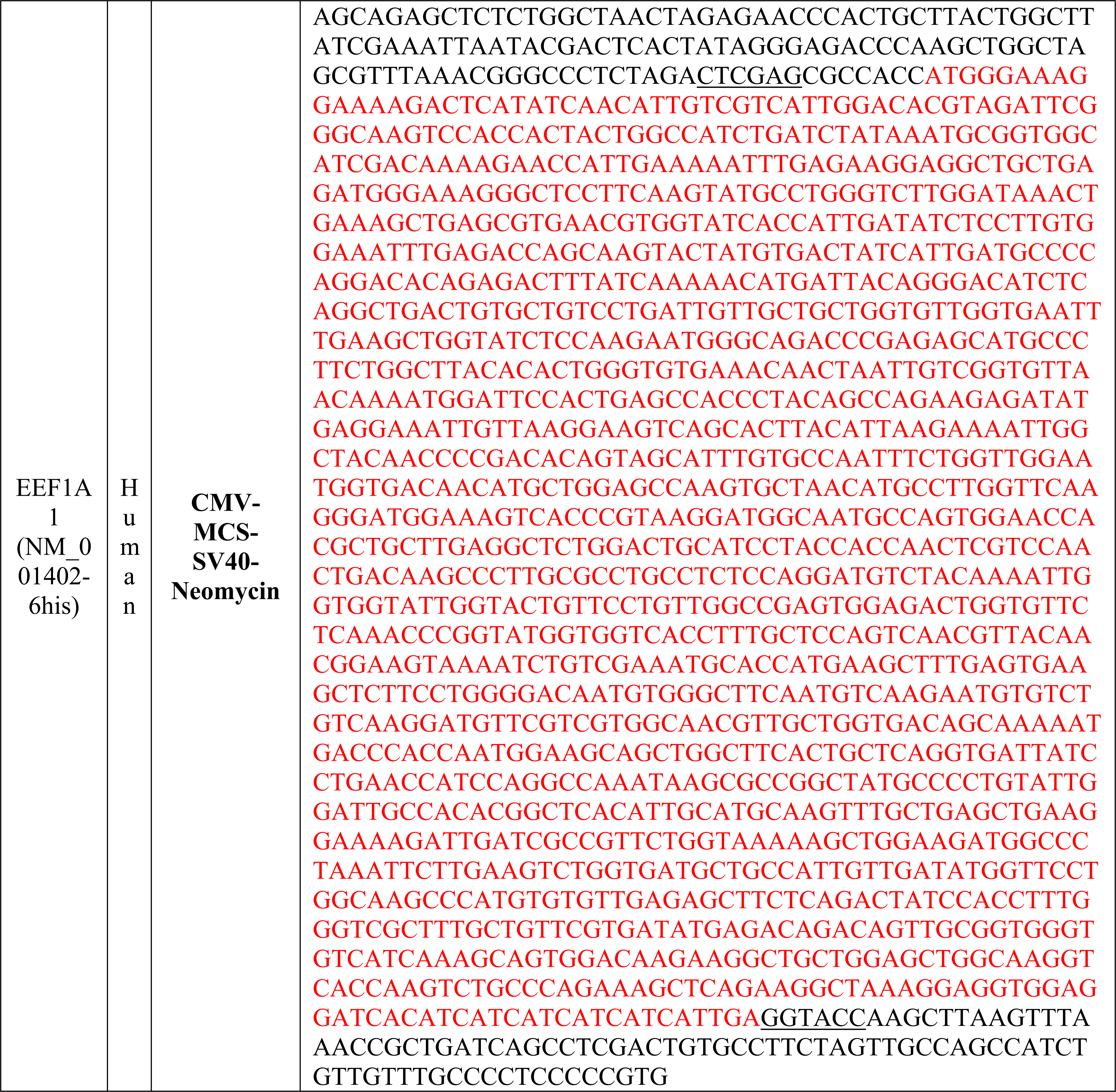
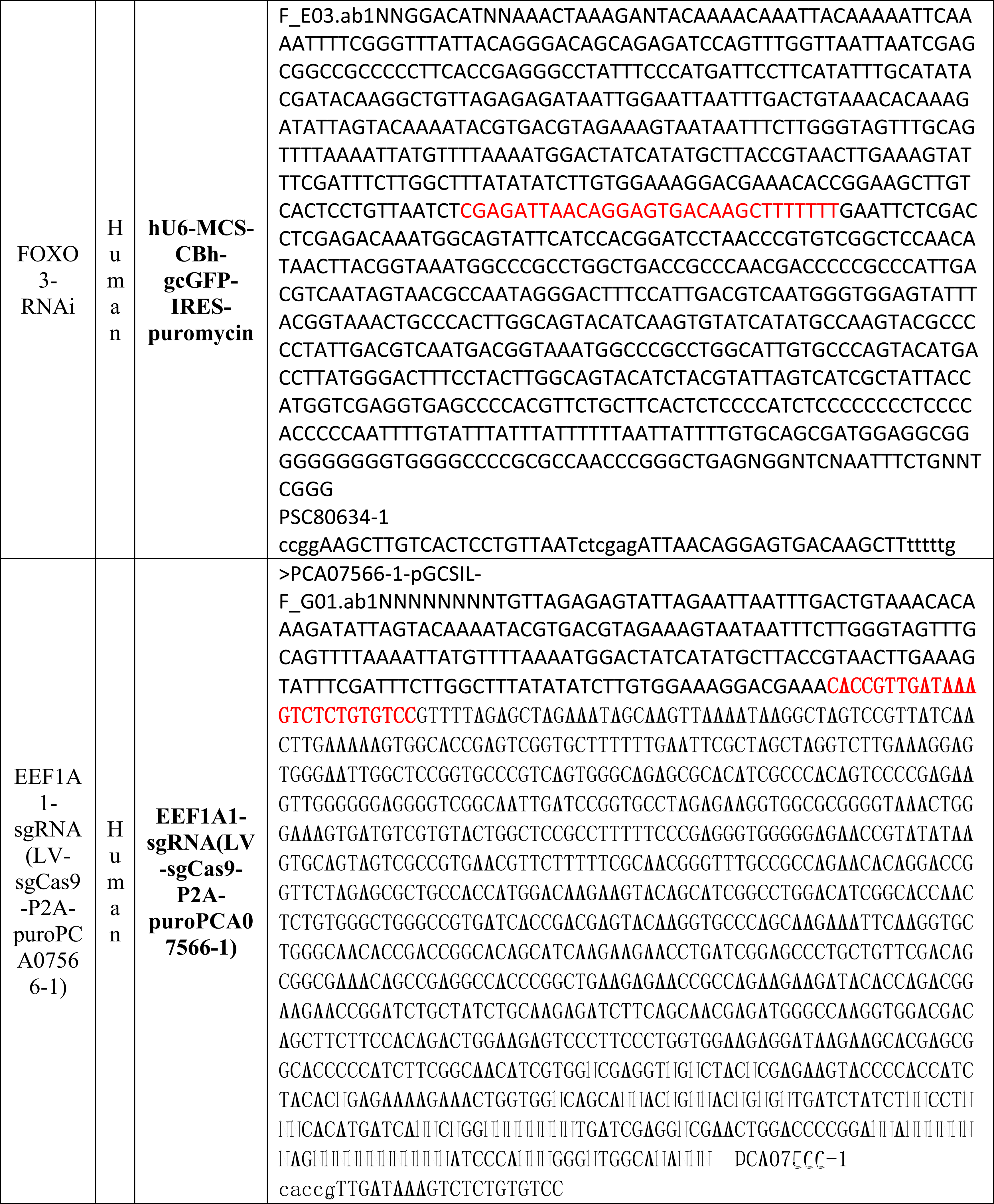
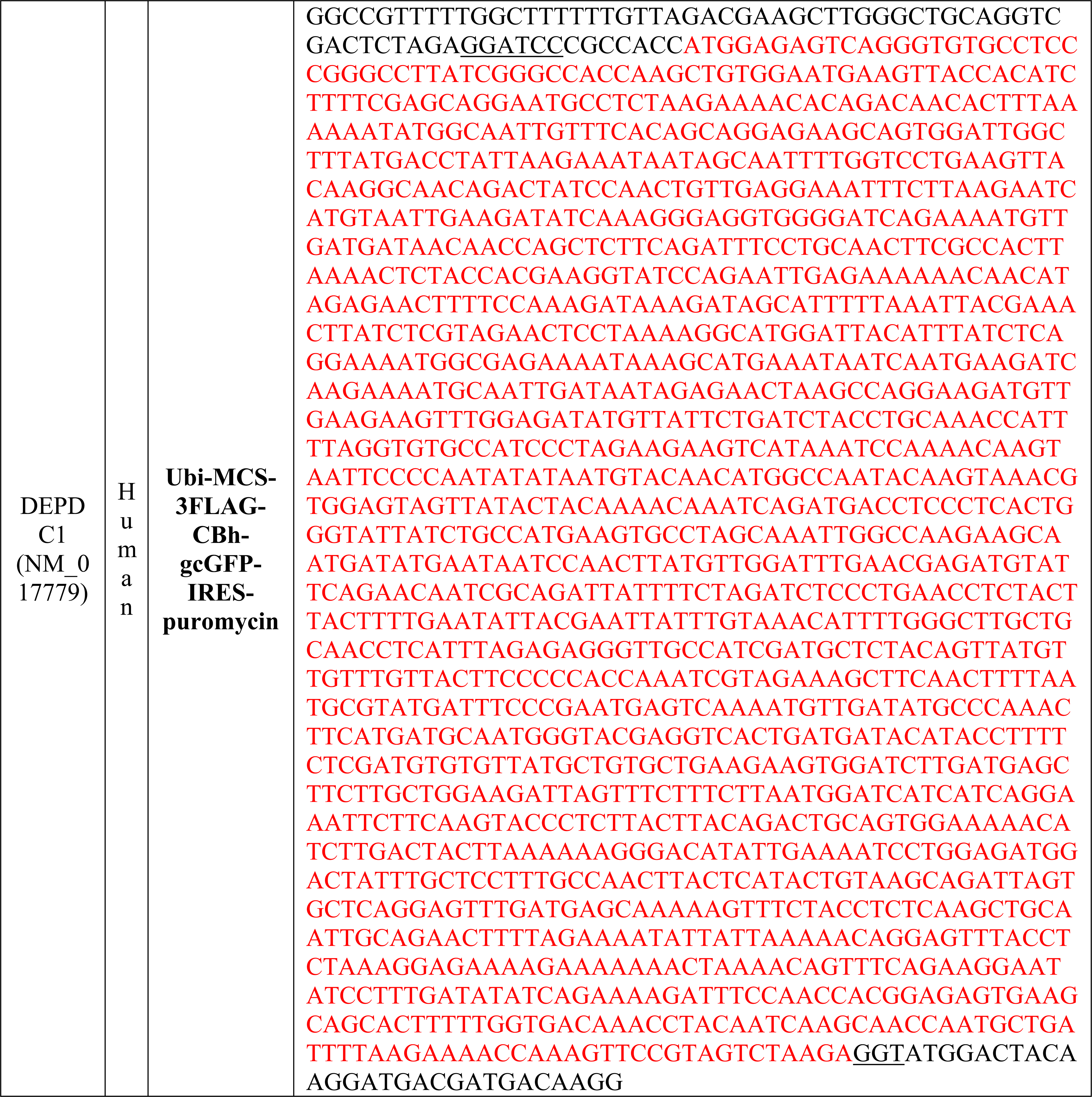
Construction of plasmids or lentivirus.

